# SWI3b restricts *AGO5* expression to promote the initiation of megagametogenesis in Arabidopsis

**DOI:** 10.64898/2026.07.26.740753

**Authors:** Wen Gong, Liping Liu, Hanyang Cai, Thomas Dresselhaus

**Affiliations:** Plant Cell Biology, Biochemistry, and Biotechnology, University of Regensburg, 93040 Regensburg, Germany; College of Life Sciences, Fujian Agriculture and Forestry University, Fuzhou 350002, China

## Abstract

Female germline development is fundamental to plant reproduction. During megasporogenesis, a somatic ovule cell differentiates into a megaspore mother cell (MMC) and undergoes meiosis to produce a single haploid megaspore while three spores degenerate. The surviving functional megaspore undergoes three rounds of mitosis during the process of megagametogenesis to produce the seven-celled female gametophyte or embryo sac. The mechanism by which the functional megaspore initiates gametogenesis remained unclear. Here, we show that the chromatin remodelling protein SWI3b is required for the initiation of megagametogenesis, but not for megasporogenesis. SWI3b activity either in the MMC or somatic ovule primordium cells alone is not sufficient for the initiation of megagametogenesis. We identified *AGO5* as a direct target of SWI3b, which restricts *AGO5* expression to the nucellus. We further show that SWI3b is required for maintaining low histone H3 lysine 9 acetylation level at the *AGO5* locus, while elevated AGO5 abundance in the nucellus disrupts the initiation of megagametogenesis. In summary, our study reveals that SWI3b promotes the initiation of megagametogenesis through chromatin-mediated repression of *AGO5*, leading to its exclusion from the MMC and thereby linking histone acetylation and epigenetic mechanisms to the development of the female germ cells.

**Highlights:**

- The chromatin remodeling complex component SWI3b is required for the initiation of megagametogenesis and early embryo development.
- In ovule primordia SWI3b restricts *AGO5* expression to the nucellus
- SWI3b acts via regulating histone acetylation around the *AGO5* transcription start site.
- Development of female germ cells are associated to histone acetylation

## Introduction

SWITCH/SUCROSE NONFERMENTING (SWI/SNF) chromatin remodeling complexes mediate ATP-dependent alterations of DNA-histone interactions and regulate DNA accessibility through nucleosome assembly, sliding, eviction, and histone variant incorporation (Clapier *et al*., 2017; Huang *et al*., 2025). SWI/SNF was first discovered through genetic screens for mating-type switch defects and sucrose non-fermenting genes in yeast (Stern *et al*., 1984). Subsequently, homologs of SWI/SNF were characterized in various eukaryotes, including Drosophila, mammals and plants (Singh *et al*., 2023). There are two SWI/SNF ATPases (Swi2 and Sth1) in yeast, one SWI/SNF ATPase BRAHMA (BRM) in Drosophila and two SWI/SNF ATPases (BRM and BRG1) in humans (Hodges *et al*., 2016). The SWI/SNF ATPases form two to three classes of multi-subunit complexes in yeast, Drosophila and mammals (Hodges *et al*., 2016; Mashtalir *et al*., 2018; Michel *et al*., 2018).

The SWI/SNF subfamily in *Arabidopsis thaliana* (Arabidopsis) includes four ATPase chromatin remodelers: BRM, SYD (SPLAYED), MINU1 (MINUSCULE 1) and MINU2. The subunits of the SWI/SNF complexes in plants were predicted based on sequence similarity and identified through forward or reverse genetic analyses (Han *et al*., 2015; Sarnowska *et al*., 2016). However, the full composition of SWI/SNF in Arabidopsis was revealed only in recent years through immunoprecipitation–mass spectrometry analyses. Three distinct subcomplexes: BAS (BRM-associated SWI/SNF), SAS (SYD-associated SWI/SNF) and MAS (MINU1/2-associated SWI/SNF) were identified (Guo *et al*., 2022; Fu *et al*., 2023). In Arabidopsis, four SWI3 paralogs have been discovered, SWI3A, SWI3B, SWI3C and SWI3D (Sarnowski *et al*., 2005). Among them, SWI3C and SWI3D were exclusively present in the BAS and SAS subcomplexes, respectively. SWI3A and SWI3B were specifically associated with the MAS subcomplex (Guo *et al*., 2022). The three subcomplexes share components, including BCL7A/B, ARP4/7 (ACTIN-RELATED PROTEIN 4/7) and SWP73A/B. Compared with mammalian SWI/SNF complexes, the MAS subcomplex evolved multiple plant-specific subunits (SHH2, PSA1 and PSA2) (Fu *et al*., 2023). The mutants of Arabidopsis SWI/SNF complexes display pleiotropic developmental defects (Huang *et al*., 2025). Studies in Arabidopsis indicated that the inactivation of MAS subunits (SWI3a, SWI3b, MINU1/2, SWP73, LFR, ARP7, PSA2) lead to lethality at reproductive stages. Among them, mutation in SWI3B, ARP7 and PSA2 showed female gametophytic defects (Kandasamy *et al*., 2005; Sarnowski *et al*., 2005; Sang *et al*., 2012; Sacharowski *et al*., 2015; Kong *et al*., 2020). In the MAS complex, SWI3b associates with HDA6 to co-repress the transcription of a subset of transposons by modulating DNA methylation, histone modifications and nucleosome occupancy (Zhu *et al*., 2013; Yang *et al*., 2020).

Female gamete development in Arabidopsis comprises two phases: megasporogenesis and megagametogenesis. During megasporogenesis, a subepidermal diploid somatic ovule cell differentiates into a megaspore mother cell (MMC). The MMC undergoes meiosis, producing four haploid megaspores of which three degenerate. During gametogenesis, the one surviving functional megaspore (FM) undergoes three rounds of mitosis to produce the female gametophyte or embryo sac, containing the female gametes, egg and central cell, which develop into embryo and endosperm after double fertilization (Dresselhaus *et al*., 2016; Hater *et al*., 2020). MMC specification has been shown to be regulated at different levels involving signaling by hormones, small RNA pathways, cell cycle regulation and other factors (Olmedo-Monfil *et al*., 2010; Schmidt *et al*., 2011; Tucker *et al*., 2012; She *et al*., 2013; Hernández-Lagana *et al*., 2016; Cai *et al*., 2017; Li *et al*., 2017; Zhao *et al*., 2017; Zhao *et al*., 2018; Pinto *et al*., 2024; Cai *et al*., 2026). It is known that differentiation of germline precursor cells encompasses extensive DNA demethylation and histone replacement (Seki *et al*., 2005). In Arabidopsis, RNA-directed DNA methylation (RdDM) pathway component NA-DEPENDENT RNA POLYMERASE6 (RDR6), SUPPRESSOR OF GENE SILENCING3 (SGS3), ARGONAUTE9 (AGO9), AGO4 and AGO6 control early megaspore formation (Olmedo-Monfil *et al*., 2010; Hernández-Lagana *et al*., 2016). In comparison with the knowledge about megasporogenesis, little is known about the pathways involved in the transition from megasporogenesis to megagametogenesis. One study showed that AGO5 is involved in the initiation of megagametogenesis in the functional megaspore, which links sRNA pathways with the transition from sporogenesis to gametogenesis (Tucker *et al*., 2012).

In this study, we show that MINU-associated SWI/SNF (MAS) complex component SWI3b is required for female germline development, specifically the initiation of megagametogenesis. *swi3* mutant passed sporogenesis but did not proceed with gametogenesis. Spatial temporal expression of SWI3b in the MMC is sufficient to complement the gametophytic defects of the mutant. Expression level of the genes linked to cell cycle progression are reduced in *swi3*. SWI3b physically interacts with another MAS complex component SHH2. Like SWI3b, SHH2 is required for megasporogenesis and exhibits a maternal effect on fertility. Furthermore, SWI3b is required for the repression of *AGO5* expression in MMC and removal of H3K9ac at the TSS of *AGO5*. These results indicate that the MAS complex inhibits sRNA pathways in female germline cell by chromatin remodeling, which contribute to gametophyte development in Arabidopsis.

## Materials and Methods

### Materials and growth conditions

*Arabidopsis thaliana* (L.) Heynh. (Arabidopsis) ecotype Columbia (Col-0), mutant, and reporter lines were grown on soil under long-day conditions (16 h light at 8500 lux, 21°C, and 65% humidity). The mutant used in this study *is swi3-2/+* (GABI_302G08). Primers used for genotyping PCR are listed in Table S1.

### Plasmid construction

Primers used for cloning are listed in Table S1. For *pSWI3b:SWI3b-GFP*, 2029 bp upstream of the *SWI3b* (AT2G33610) start codon was taken as the promoter. The whole genomic sequence without the stop codon was amplified from genomic DNA. For *pABI3:SWI3*, 4962 bp upstream of the *ABI3* start codon was used as the promoter and the SWI3b coding sequence (CDS) was amplified from cDNA generated from Col-0 inflorescence. For *pKNU:SWI3b-GFP*, 2001 bp upstream of *KNU* was taken as the promoter. For *pWUS:SWI3-GFP*, 3439 bp upstream of the *WUS* start codon was taken as the promoter. For *pAGO5:NLS-2xGFP*, 3128 bp upstream of the *AGO5* start codon was taken as the promoter. For all GFP fusion constructs, a GSAGAG-linker was added between the CDS and the *GFP* sequence. For *pKNU:AGO5* and *pSWI3b:AGO5*, *AGO5* CDS was amplified from Col-0 inflorescence cDNA. Amplified fragments were assembled using InFusion (Takara) cloning to pGreenII0125 containing a *35S* terminator. The resistance is kanamycin in *Escherichia coli* and norflurazon *in planta*.

### Microscopy

Phenotypic analysis of ovule and embryo development was made by using differential interference contrast (DIC) microscopy (Zeiss, Imager.A2). Flower buds and siliques at different developmental stages were dissected, followed by transferring of ovules connected by the septum to a drop of clearing solution (chloral-hydrate: H₂O:glycerol = 8 : 2 : 1 solution). After the tissues were cleared, the samples were observed under the DIC microscope using a 63x objective. Expression pattern of reporter lines was observed using a spinning - disk confocal laser microscope system (Visitron system VisiScope with CSU-W1) equipped with an HC PL APO 63×/1.40-0.60 oil objective. Venus and YFP were excited at 505 nm, aniline blue staining with DAPI was excited at 405 nm, and GFP was excited at 488 nm.

### Callose detection

Callose detection in megaspores was performed as previously described (Huang *et al*., 2023). The inflorescences were fixed in formaldehyde/alcohol/acetic acid for 16 h and incubated in 0.1% (w/v) aniline blue in 50 mM phosphate buffer (pH 11) for 8–12 h. Stained pistils were mounted in 10% (v/v) glycerol and observed with the DIC microscopy (Zeiss, Imager.A2).

### RT-qPCR

For Fig. 3e, RNA samples were extracted from flowers. For Fig. 5d, RNA samples were extracted from ovules at pre-meiotic stage attached to the septum. qPCR was performed with gene-specific primers listed in Table S1 using the Eppendorf ep realplex Mastercycler real-time PCR system and the SYBR solution (Roche) as described previously and *PP2AA3* (AT1G13320) as an internal control gene (Ferreira *et al*., 2023). Three biological replicates and three technical replicates for each sample were performed in qPCR experiments.

### Chromatin immunoprecipitation

For each chromatin immunoprecipitation (ChIP) experiment, 1.5 g of inflorescence was used. Collected tissues were cross-linked using formaldehyde. Then the chromatin was extracted and sheared as previously described (Gendrel *et al*., 2005). Immunoprecipitation was performed by using specific antibodies including a polyclonal GFP antibody (ab290; Abcam), anti-H3K9ac (07-352, Millipore), and anti-H3K14ac (ABP56691, Abbkine). Relative enrichment of associated DNA fragments was analyzed by qPCR. All primers used are listed in Table S1. Three biological replicates were performed, and three technical repeats were carried out for each biological point. The presented data are from one representative experiment.

## Results

### SWI3b is required for female gametogenesis and embryogenesis

A previous study identified two T-DNA insertion mutant alleles in the gene *SWI3b*, named *swi3b-1* and *swi3b-2*. Both alleles were homozygous embryo lethal and exhibited gametophytic defects (Sarnowski *et al*., 2005). To determine the stage of gametogenesis at which these defects arise, we obtained the T-DNA mutant *swi3b-2* (hereafter referred to as *swi3-2*) for further analysis (Fig. S1). To test whether SWI3b plays a role in MMC specification, we performed DIC microscopy of premeiotic ovules. In heterozygous *swi3-2/+* mutant, 93.5±1.4% (n=310) of ovules contained one enlarged MMC, which is comparable with WT ovules (94.9±1.1%, n=305) (Fig. 1a). Since female gametophyte stage 7 (FG7) is the final, mature stage of megagametogenesis in Arabidopsis, representing the functional female gametophyte for fertilization (Skinner & Sundaresan, 2018; Hater *et al*., 2020), we next analyzed the ovules at FG7 stage. 97.5±1.1% (n=187) of WT ovules contained an egg cell, two synergids and one central cell nuclei in the embryo sac. However, 51.6±1.5% (n=197) of *swi3/+* ovules contained a one-nucleate female gametophyte similar to FG1 stage of WT ovules (Fig. 1b), suggesting that FG development had terminated prior to the first mitotic division. Furthermore, 23.9±0.7% of segregating embryos from heterozygous *swi3/+* mutant exhibited abnormal cell division patterns (anticlinal cell division in embryo proper and hypophysis) from 8-cell to 16-cell embryo stage, instead of the periclinal cell division to form the protoderm and asymmetric cell division of the hypophysis in WT (Fig. 1c, 1d). Embryos with abnormal patterning arrested later at the globular stage (Fig. S1). To investigate the inheritance of the arrested ovule phenotype, we performed reciprocal crosses between heterozygous swi3-2/+ and WT plants. When WT pollen was used to pollinate *swi3-2/+* pistils, we observed a ∼1:1 segregation of fertilized and unfertilized ovules in the F1 seeds, which is consistent with the segregation ratio of a female gametophytic defect, comparable to that of *swi3-2/+* pollen used to pollinate *swi3-2/+* pistils. When *swi3-2/+* pollen was used to pollinate WT pistils, we observed nearly full seed set, similar to that obtained with WT pollen (Fig. 1f, 1g). Next, we examined the expression pattern of *pSWI3b:SWI3b-GFP* during ovule development and embryogenesis. SWI3b-GFP was observed in both the MMC and somatic cells in the premeiotic ovule and remained ubiquitously expressed after meiosis from FG1 to FG7 (Fig. 1e, Fig. S2). After fertilization, SWI3b-GFP was observed throughout embryogenesis, showing ubiquitous expression until the globular stage and becoming concentrated in the epidermis from the late globular stage onwards (Fig. 1e). *pSWI3b:SWI3b-GFP* complemented both, the gametophytic and embryo defects of the mutant (Fig. S3a-d), indicating that the fusion protein is functional.

**Fig. 1.**
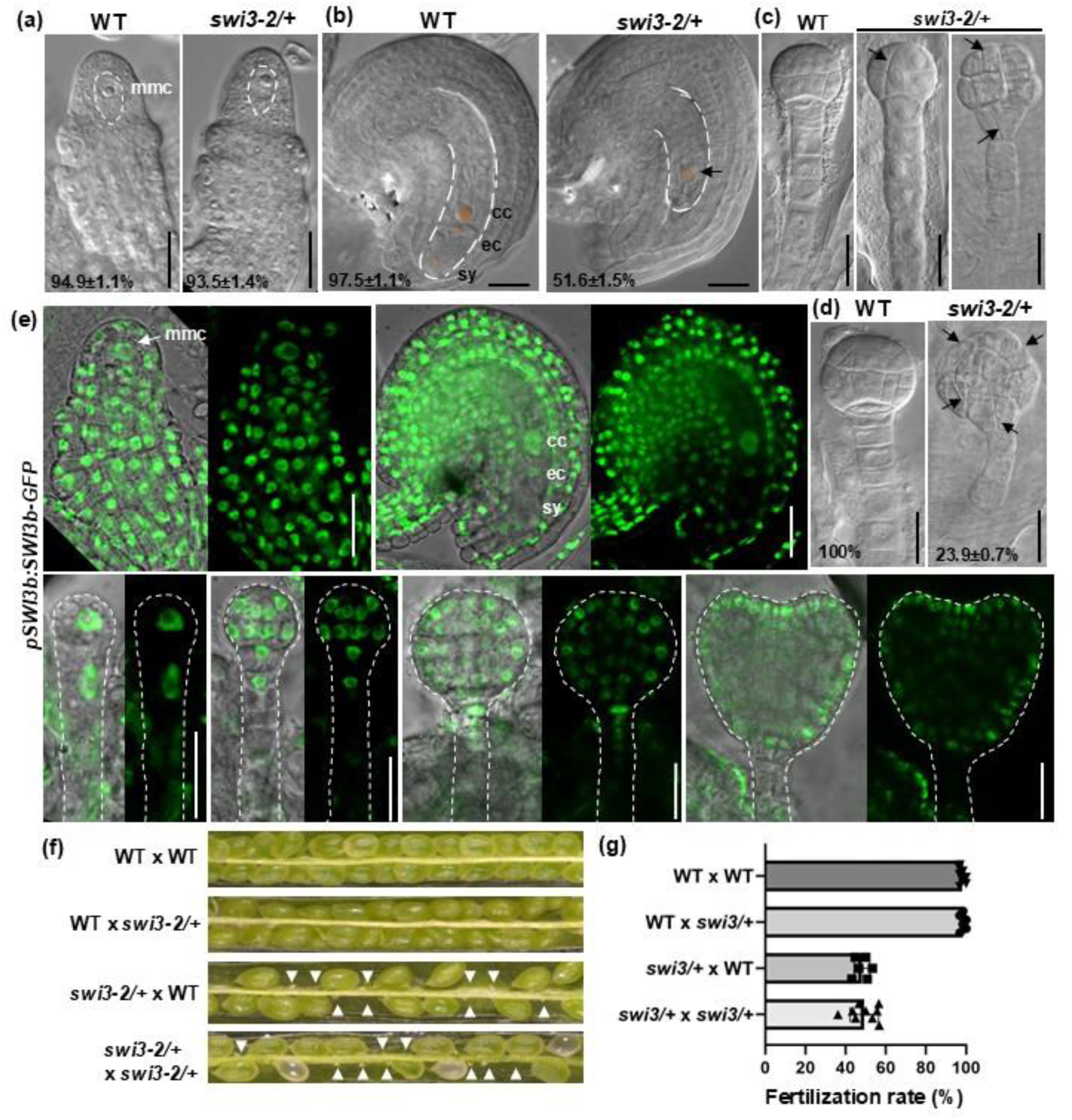
SWI3b is required for female gametogenesis and embryo development. (a) DIC microscopy of premeiotic ovules in WT and *swi3-2/+*. Dashed line indicates the area of the megaspore mother cell (MMC). Bars, 10 μm. (b) DIC microscopy of ovules at FG7 in WT and *swi3-2/+*. Dashed line indicates the embryo sac. Arrow points to the single nucleus. Bars, 20 μm. (c-d) DIC microscopy of 8-cell to 16-cell stage (c) and globular stage (d) embryos in WT and *swi3-2/+*. Arrows point to abnormal cell division planes. Bars, 20 μm. (e) Expression pattern of *pSWI3b:SWI3b-GFP* in premeiotic ovules and mature ovules at FG7 (upper row) and embryos from 1-cell stage to heart stage (lower row). White dashed lines indicate the outline of the embryo. Bars, 20 μm. (f) Dissected siliques from the reciprocal crosses between WT and *swi3-2/+*. White arrowheads indicate aborted ovules. (g) Quantification of the fertilization rate from reciprocal crosses.

### SWI3b is required for the initiation of megagametogenesis

To confirm that megasporogenesis is not affected by mutation of *SWI3b*, we examined the expression of *pKNU:KNU-Venus*, a marker line for MMC identity (Su *et al*., 2020), in WT and *swi3-2/+*. KNU-Venus was detected in a single MMC in 93.9% (n=308) of pre-meiotic ovules in *swi3-2/+*, comparable to 95.4% (n=328) in WT ovules (Fig. 2a), suggesting that megasporogenesis was normal in the mutant. Next, we investigated whether the MMC of the mutant undergoes meiosis using aniline blue staining to detect callose deposition. Callose deposition was observed in 95.4±0.7% (n=345) *swi3-2/+* ovules, which is comparable to 91.7±0.2% in WT ovules (Fig. 2), suggesting that mutant MMCs are capable to enter meiosis. After meiosis, the surviving functional megaspore (FM) undergoes three rounds of mitosis to produce the female gametophyte. To investigate whether the mutant FM enters mitosis, we analyzed the expression of the gametophytic cell marker *pAKV:H2B-YFP* (Yang *et al*., 2010). *pAKV:H2B-YFP* was detected in the FM of both, WT and mutant ovules at FG1, indicating that FM formation occurs normally in the mutant. In WT ovules, *pAKV:H2B-YFP* was subsequently observed in multiple gametophytic nuclei. In contrast, in 47.2±1.9% of *swi3-2/+* ovules (n=311), *pAKV:H2B-YFP* was detected in only a single nucleus (Fig. 2c), suggesting that FG1 abortion in *swi3-2/+* is caused by defective mitosis. After three rounds of mitosis, the FM gives rise to a mature female gametophyte containing an egg cell and a central cell, both of which are destined for fertilization. To further determine whether the single nucleus in *swi3-2/+* at FG7 acquires generative cell identity, we analyzed the egg cell marker *pEC1:EC1-GFP* and the central cell marker *pDD65:NLS-GFP*. In about half of the *swi3-2/+* ovules, expression of neither of the two markers was detected (Fig. 2d) indicating that the single nucleus in *swi3-2/+* did not acquire generative cell identity.

**Fig. 2.**
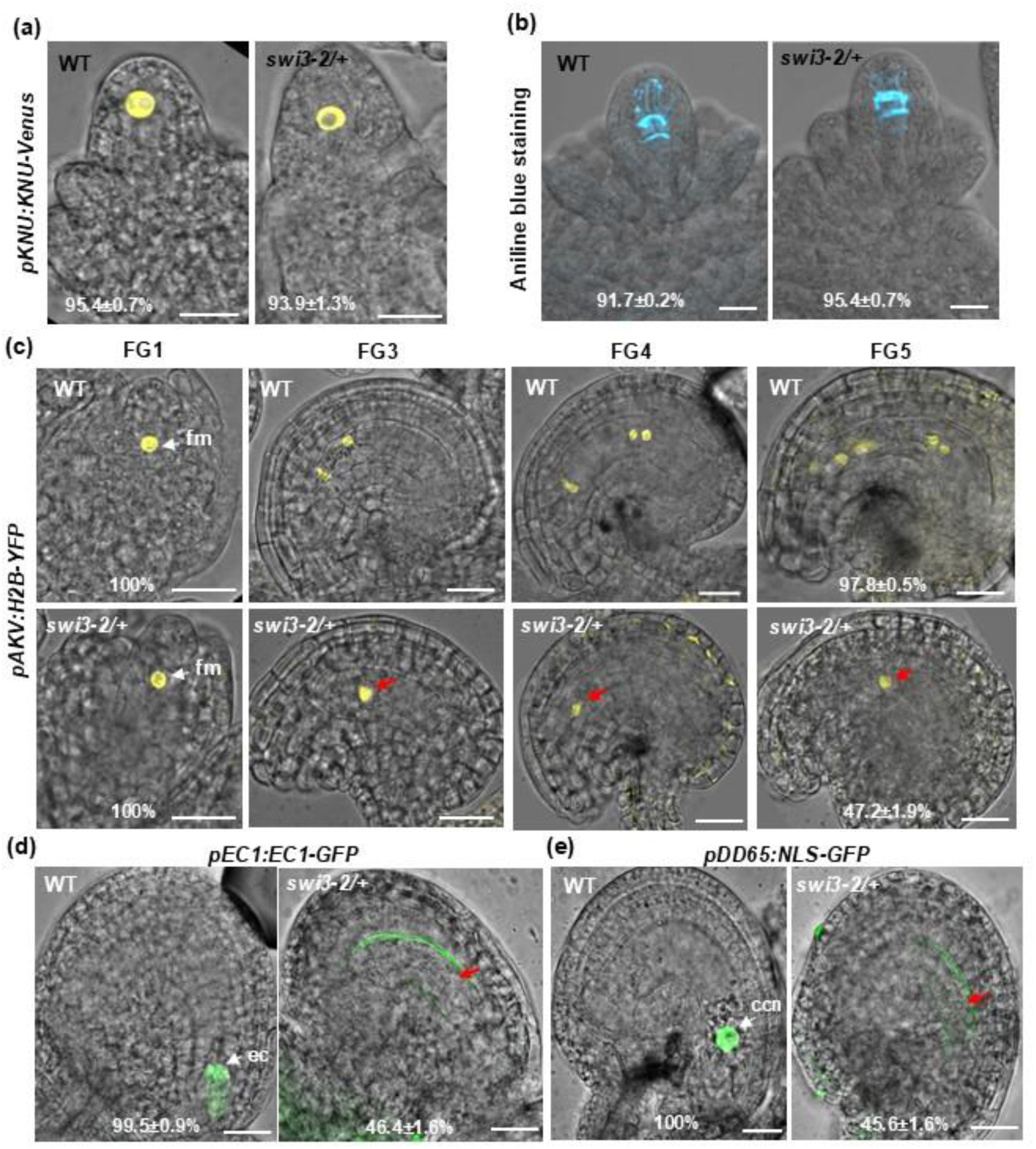
SWI3 is required for mitotic divisions of the functional megaspore. (a) Expression pattern of *pKNU:KNU-Venus* in premeiotic ovules of WT and *swi3-2/+*. (b) Aniline blue staining showing callose deposition in WT and *swi3-2/+* ovules at meiosis stage. (c) Expression pattern of *pAKV:H2B-YFP* in WT and *swi3-2/+* ovules from FG1 to FG5. (d) Expression pattern of *pEC1:EC1-GFP* in WT and *swi3-2/+* ovules at FG7. (e) Expression pattern of *pDD65:NLS-GFP* in WT and *swi3-2/+* ovules at FG7. Red arrows indicate the single nucleus. Abbreviations: ccn: central cell nuclei; ec: egg cell; fm: female gametophyte. Bars, 10 μm.

### Embryo rescued *swi3* showed defects in the initiation of megagametogenesis and reduced expression of *CDKA*

Given that mutation in *SWI3b* showed both, female gametophytic and embryonic defects and is homozygous embryonic lethal, we sought to generate a homozygous *swi3* line with the expression of *SWI3b* during embryo development, but which limits the production of SWI3b beyond this stage. The expression of the *ABI3* gene has been reported previously, and its promoter has been used to rescue other embryo-lethal mutation to study vegetative gene function (Despres *et al*., 2001; Bodi *et al*., 2012). The *SWI3b* coding sequence, expressed under the control of the *ABI3* promoter, was introduced into *swi3-2/+* plants. Two independent transgenic lines (*pABI3:SWI3b swi3-2/+*) were analyzed. The frequency of embryo defects was reduced to 16.6±4.3% and 6.1±3.1% in T2 line 1 and line 2, respectively, compared with 24.1±1.5% in *swi3-2/+* (Fig. S4a, b). Among the progeny, homozygous *swi3-2* plants were identified, indicating that this construct was able to complement the embryo-lethal phenotype. *pABI3:SWI3b swi3-2* (hereafter referred to as *ABI3 swi3-2*) homozygous plants displayed various developmental defects, including smaller rosette leaves, shorter stems, and reduced branching (Fig. S4c). This indicates that SWI3b plays roles in multiple developmental processes. We found that expression levels of *SWI3b* in *ABI3 swi3-2* were indeed generally significantly reduced compared with that of WT plants. The reductions were more pronounced in flowers and inflorescences, showing approximately 3-fold and 2-fold decreases, respectively (Fig. S4d). In homozygous *ABI3 swi3-2* plants, 99.5±0.9% of embryos showed normal patterning, similar to WT plants (Fig. 3a). By contrast, 58.4±10.4% of *ABI3 swi3-2* ovules at the FG7 stage contained a single nucleus, indicating loss of SWI3b function during female gametogenesis (Fig. 3b). Reciprocal crosses between *swi3-2/+* and WT revealed that the fertilization defect arises from a female gametophytic defect (Fig. 1f-g). However, using *swi3-2/+* as the pollen doner may result in excess WT pollen masking any effect of *swi3-2* pollen. Consistent with pollen transcriptome data showing that *ABI3* is not expressed in pollen (Loraine *et al*., 2013), we performed reciprocal crosses between *ABI3 swi3* and WT to assess potential pollen contributions. When WT pollen was used to pollinate *ABI3 swi3-2* pistils, we observed a ∼1:1 segregation of fertilized and unfertilized ovules, consistent with a female gametophytic defect and comparable to crosses using *ABI3 swi3* pollen on *ABI3 swi3-2* pistils. In contrast, *ABI3 swi3* pollen fertilized WT pistils efficiently, producing nearly full seed set, similar to WT crosses (Fig. 3c-d). Taken together, these results indicate that SWI3b is required for female gametogenesis but is largely dispensable for male gametogenesis.

**Fig. 3.**
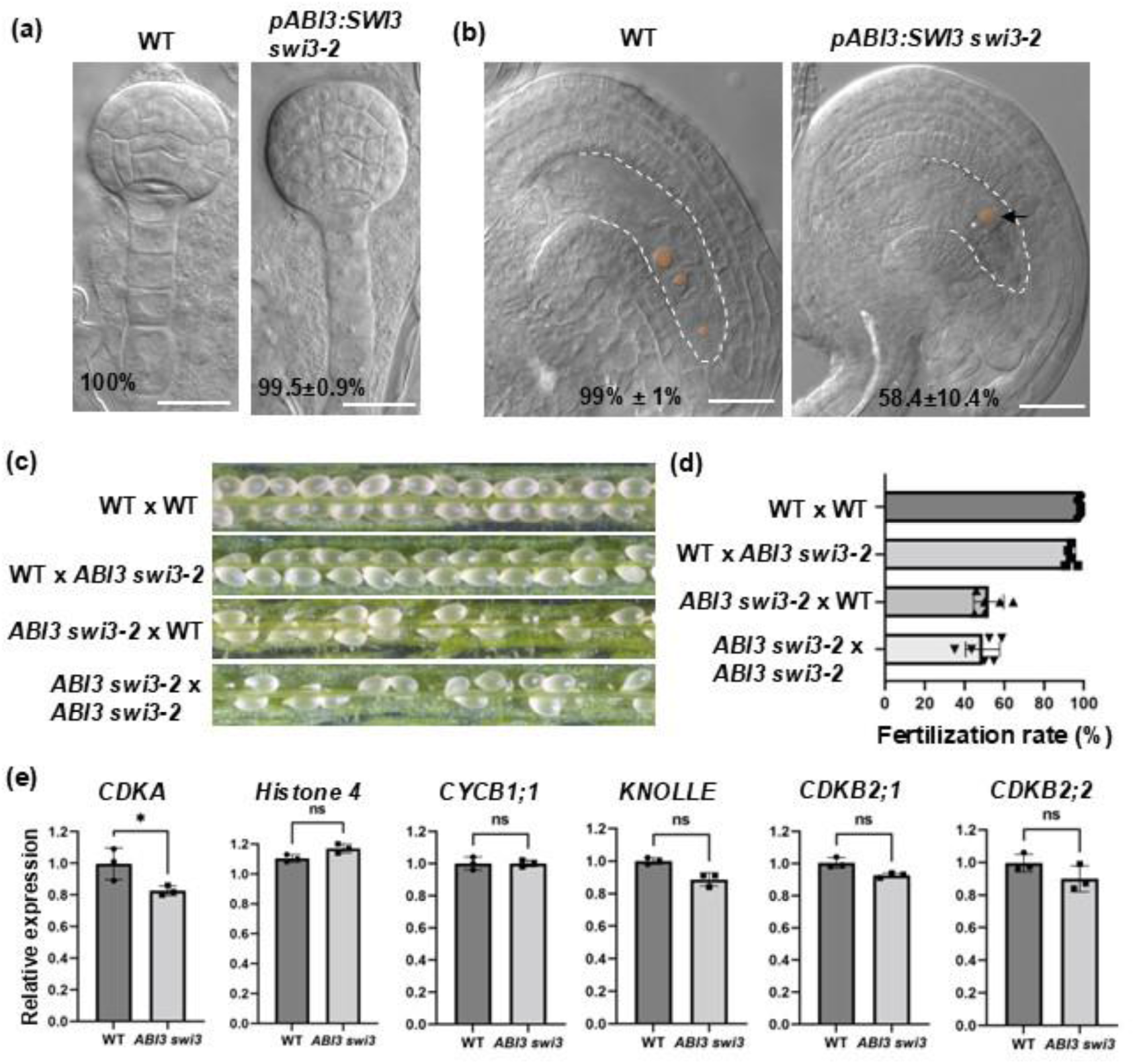
Embryo rescued *swi3/-*mutants showed defects in megagametogenesis and reduced expression of *CDKA*. (a) DIC microscopy of ovules at FG7 in *swi3-2/+* and *ABI3 swi3-2*. Dashed line indicates the embryo sac. Arrow points to the single nucleus. Bars, 20 μm. (b) DIC microscopy of globular embryo stage in *swi3-2/+* and *ABI3 swi3-2*. Arrows point towards abnormal cell division planes. Bars, 20 μm. (c) Dissected siliques from the reciprocal crosses between WT and *ABI3 swi3-2*. (g) Seed numbers from reciprocal crosses. (e) Transcripts level of cell cycle regulator genes in WT and *ABI3 swi3-2* pistils, determined by quantitative RT-PCR. Error bars indicate SD (n = 3). Unpaired two-tailed Student’s t-test is employed to measure statistical significance between two samples (ns, no significance; *, *p* < 0.05).

Since mitosis of the FM in *swi3* was compromised, we hypothesized that expression of genes regulating the cell cycle are mis-expressed. To test this hypothesis, we examined the expression of genes linked to cell cycle progression in WT and *ABI3 swi3-2* pistils. *CDKA* is a general regulator of cell cycle progression whose activity peaks at the G1/S and G2/M transitions. *Histone H4* and *KNOLLE* are specifically expressed in S-and M-phase, respectively; *CYCB1;1* is a mitotic cyclin required for G2/M transition of the cell cycle, while *CDKB2;1* and *CDKB2;2* are expressed at the G2/M boundary (Vandepoele *et al*., 2002). Significant differences were not found in the expression level of *Histone 4*, *CYCB1;1*, *KNOLLE*, *CDKB2;1* and *CDKB2;2*. However, a significant reduction in the expression of *CDKA* was found in *ABI3 swi3-2*, compared with WT plants (Fig. 3e). This is consistent with previously reported functions of CDKA in megagametogenesis (Dissmeyer *et al*., 2009).

### SWI3b activity in the MMC or epidermal cells alone was not sufficient for the initiation of megagametogenesis

To determine whether SWI3b activity is required in the MMC or in the surrounding somatic nucellar cells during female gametogenesis, we expressed *SWI3b-GFP* under the control of the MMC-specific *KNUCKLES* (*KNU*) promoter and the somatic nucellar epidermis-specific *WUSCHEL* (*WUS*) promoter in the *swi3-2/+* background. *pKNU:SWI3b-GFP* was detected in the MMC of pre-meiotic ovules, with weaker signal at the neighbouring cells (Fig. 4a), which was slightly different from the MMC-specific expression of *pKNU:KNU-GFP* (Fig. 2a). In ovules at FG1, *pKNU:SWI3b-GFP* was detected in the FM and neighboring cells (Fig. 4b). *pWUS:SWI3b-GFP* was detected in somatic nucellar epidermal cells of pre-meiotic ovules (Fig. 4c) and then more broadly in the whole nucellus at FG1 (Fig. 4d). Expression of *SWI3b-GFP* under control of the *KNU* promoter showed about 5% restoration of normal female gametogenesis and fertilization in one transgenic line in *swi3-2/+*, and no significant difference in another line (Fig. 4e-g), suggesting that SWI3b activity in MMC alone was not sufficient for the initiation of megagametogenesis. A similar result was observed in *pWUS:SWI3b-GFP swi3-2/+*, which showed about 5% restoration of normal female gametogenesis and fertilization (Fig. 4e-g). This finding indicates that SWI3b activity in nucellar epidermis cells alone was not sufficient for the initiation of megagametogenesis. By contrast, expression of *SWI3b-GFP* by the *SWI3b* promoter complemented both, mutant effects in female gametogenesis and embryogenesis (Fig. 4 e-g). Taken together, we conclude that SWI3b activity in both MMC and nucellar somatic cells is required for proper female gametogenesis, while SWI3b activity in the MMC or somatic cells alone is not sufficient for the initiation of megagametogenesis.

**Fig. 4.**
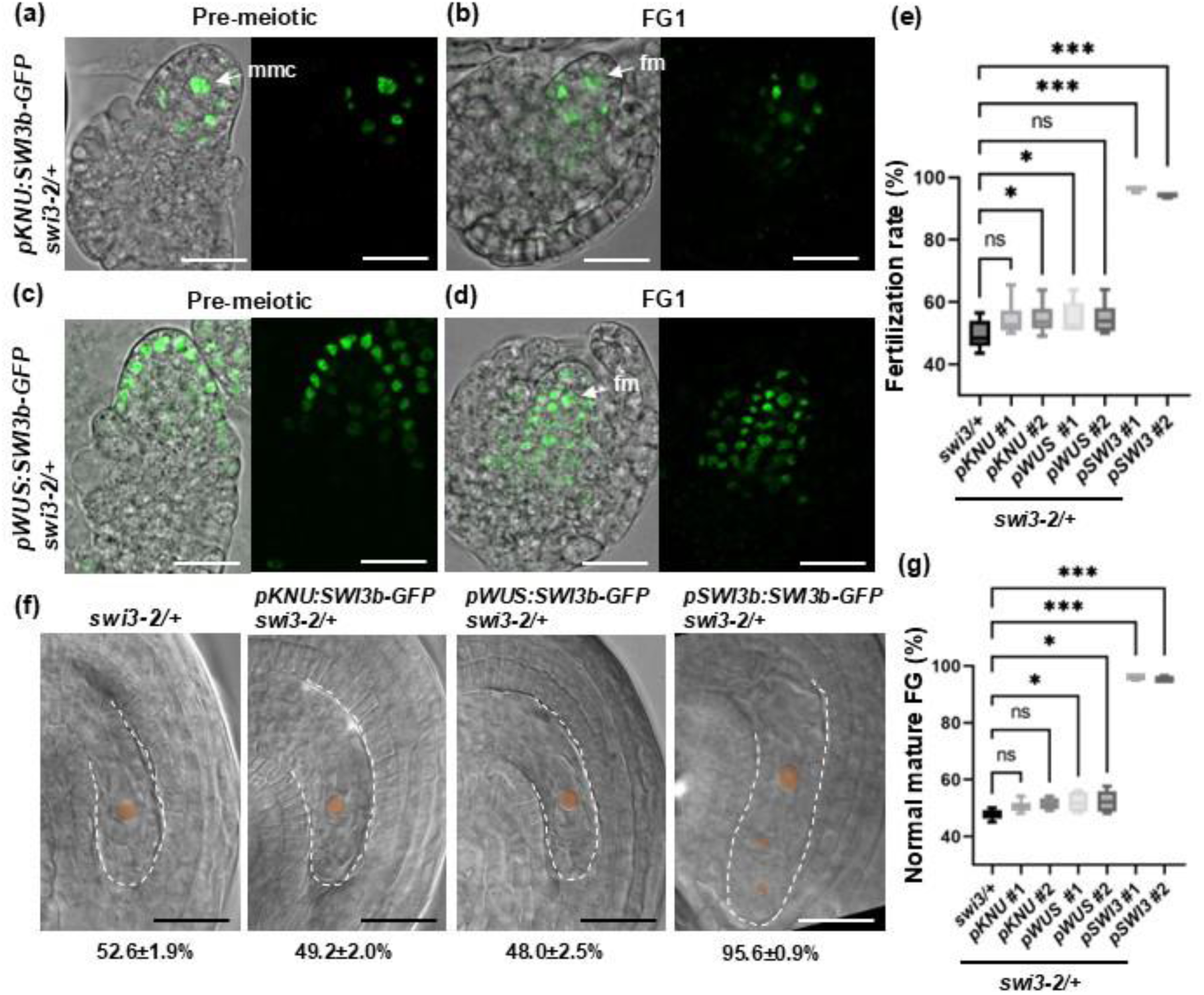
SWI3b activity in the MMC or epidermal cells of ovule primordia alone was not sufficient for proper female gametogenesis. (a-b) Expression pattern of *pKNU:SWI3b-GFP* in *swi3-2/+* at pre-meiotic stage (a) and at FG1 (b). (c-d) Expression pattern of *pWUS:SWI3b-GFP* in *swi3-2/+* at pre-meiotic stage (c) and at FG1 (d). Bars, 20 μm. (e) Box blot showing the fertilization rate of *pKNU:SWI3b-GFP* and *pWUS:SWI3b-GFP* in *swi3-2/+*. (f) DIC microscopy of ovules at FG7 in *swi3-2/+* and indicated transgenic lines in *swi3-2/+*. Dashed line indicates the embryo sac. Nuclei in the female gametophyte are marked by red color. Bars, 20 μm. (g) Box blot showing the percentage of WT-like female gametophyte of the indicated transgenic lines in *swi3-2/+*. Statistical significance was determined using one-way ANOVA followed by Dunnett’s multiple comparisons test, comparing each transgenic line with the *swi3-2/+* (ns, no significance; *, *p* < 0.05, ***, *p* < 0.001).

### SWI3b represses the expression of *AGO5* in the pre-meiotic ovule by regulating histone H3 lysine 9 acetylation

Next, we asked which downstream factor(s) of SWI3b mediate its function in regulating megagametogenesis. Previous studies have shown that ARGONAUTE5 (AGO5), an effector of small RNA (sRNA)-mediated silencing pathways, might play a role in megasporogenesis. A dominant insertion allele *ago5-4* showed defects in the initiation of megagametogenesis that resemble those observed in *swi3-2* (Tucker *et al*., 2012; Roussin-Leveillee *et al*., 2020). Because SWI/SNF complexes are recruited to acetylated nucleosomes and interact with histone deacetylases (HDACs), they have been proposed to function in histone deacetylation and transcriptional repression (Mizutani *et al*., 2002; Yang *et al*., 2020; Ramirez-Prado & Benhamed, 2021). Therefore, we hypothesized that SWI3b and its associated SWI/SNF complex regulate megagemetogenesis by restricting *AGO5* expression in the nucellus. In WT ovule primordia, the *pAGO5:NLS-2xGFP* construct showed GFP expression in the nucellar epidermis and inner integument primordia and was absent from the MMC (Fig. 5a), which is consistent with previous findings (Tucker *et al*., 2012; Pinto *et al*., 2024). Notably, in *swi3-2/+*, GFP fluorescence was detected in the MMC in about half of pre-meiotic ovules. In *ABI3 swi3-2*, GFP fluorescence was also detected in the MMC, in addition to the epidermis and inner cell layers (Fig. 5a). We next quantified the average fluorescent intensity in the MMC and across the entire nucellar region. In the MMC, the average fluorescence intensity was significantly higher in *swi3-2/+* and *ABI3 swi3-2* compared to the WT (Fig. 5b). Across the entire nucellar region, the average GFP fluorescence intensity was significantly higher in *ABI3 swi3-2* than in the WT, a slight increase was also observed in *swi3-2/+* compared with the WT (Fig. 5c). This indicates that *AGO5* expression in nucellus was repressed by SWI3b. Following meiosis at the FG1 stage, GFP fluorescent intensity remained significantly higher in the nucellus of both *swi3-2/+* and *ABI3 swi3-2* than in the WT (Fig. S5a-b). In addition, qRT-PCR analysis showed that the transcript level of *AGO5* in ovule primordia was significantly increased in *ABI3 swi3-2* compared with the WT (Fig. 5d), which is consistent with repression of *AGO5* expression by SWI3b in the ovule. To examine whether *AGO5* is a direct target of SWI3b, chromatin immunoprecipitation (ChIP)-qPCR was performed using *pSWI3b:SWI3b-GFP* inflorescence. An anti-GFP antibody was used for immunoprecipitation. The occupancy of SWI3b to the genomic regions 300 base pairs (bp) upstream of the transcriptional start site (TSS), direct after the TSS and in the first exon was analyzed (Fig. 5e). A significant enrichment of the fragments upstream of the TSS and directly after the TSS was detected in *pSWI3b:SWI3b-GFP* compared with WT plants. In contrast, significant enrichment was not detected in the region of the first exon (Fig. 5f). These data suggest that SWI3b specifically binds to the regions near the TSS of *AGO5*. To investigate whether the repression of *AGO5* expression was caused by histone deacetylation, we next determined histone H3 acetylation levels of those three regions of *AGO5* by ChIP-qPCR using antibodies specific for acetylated histone H3 at lysine 9 (K9) and lysine 14 (K14) in WT and *ABI3 swi3* inflorescences. The H3K9ac level increased in the regions upstream of the TSS and directly after the TSS in *ABI3 swi3* compared with WT inflorescences (Fig. 5g). In contrast, the H3K14ac level was not significantly changed in all three regions (Fig. 5h). As a negative control, significant differences were not detected in all three regions when antibodies were not used during immunoprecipitation (Fig. S6). Collectively, these results suggest that SWI3b directly binds to *AGO5* near the TSS and represses the expression of *AGO5* by decreasing the levels of histone H3K9ac.

**Fig. 5.**
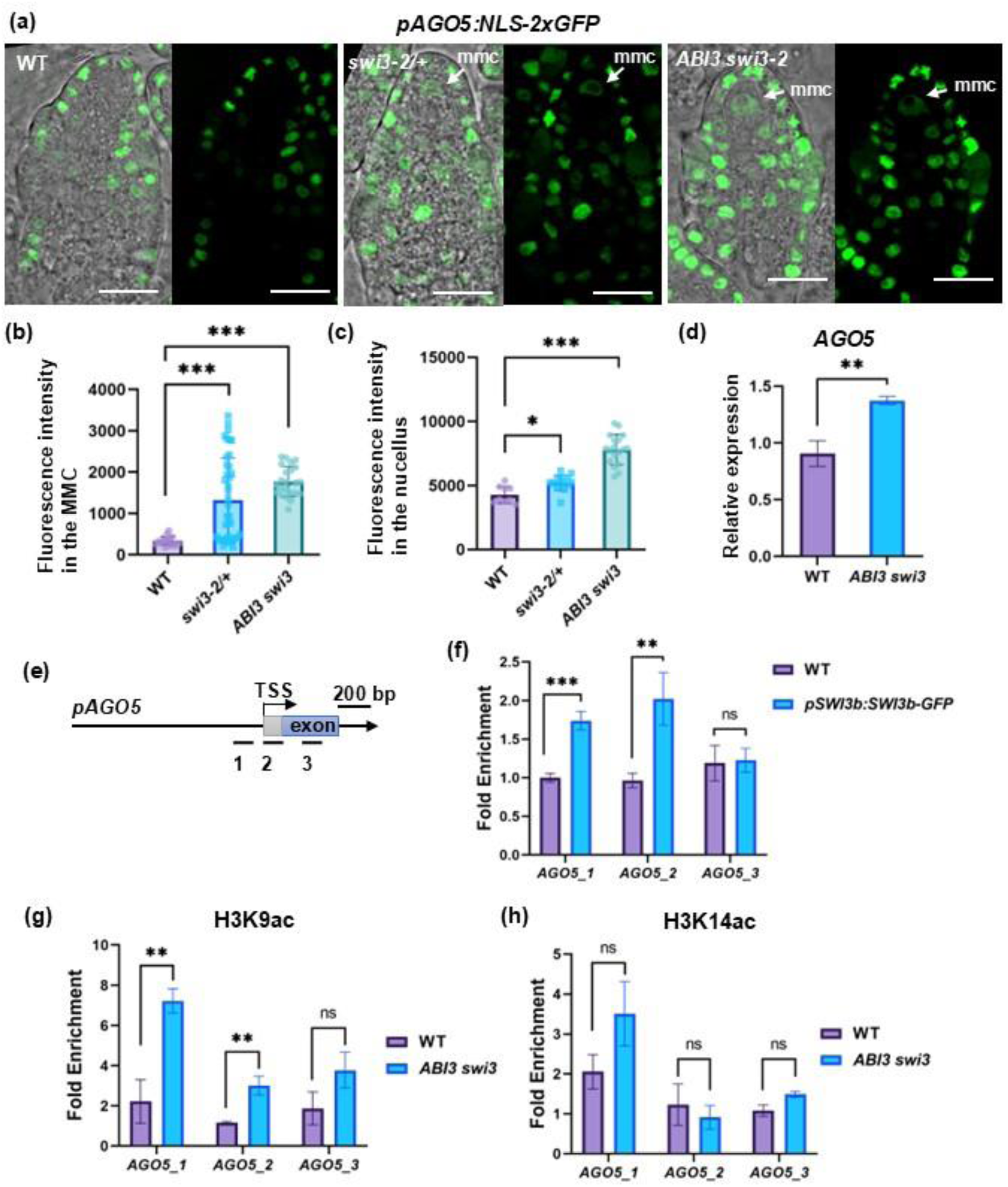
SWI3b represses the expression of *AGO5* in the pre-meiotic ovule and regulates histone H3 lysine 9 acetylation levels at *AGO5.* (a) Expression pattern of *pAGO5:NLS-2xGFP* in WT (left), in *swi3-2/+* (middle) and in *ABI3 swi3-2* (right) at pre-meiotic stage. Arrows point to the MMC. Bars, 20 μm. (b-c) Quantification of the average fluorescence intensity in the MMC (b) and in the nucellus (c) of the indicated genotypes. One-way ANOVA followed by Dunnett’s multiple comparisons test, comparing each mutant line with the WT (ns, no significance; *, p<0.05; ***, *p* < 0.001). (d) Transcripts level of cell cycle regulator genes in WT and *ABI3 swi3-2* ovules, determined by quantitative RT-PCR. Error bars indicate SD (n = 3). (e) Schematic drawing of the genomic region of *AGO5* until the first exon. The three black lines underneath indicate the fragments analyzed by qPCR. (f) ChIP-qPCR analysis showing enrichment in region 1 and 2 of *AGO5* in the inflorescence of *pSWI3b:SWI3-GFP* transgenic plants using an anti-GFP antibody. Values are means ± SD from three biological replicates. Each biological replicate corresponds to three technical replicates. (g-h) ChIP-qPCR analysis showing enrichment for H3K9ac in region 1 and 2 of *AGO5* in *ABI3 swi3-2* inflorescence compared with WT (g), and lack of enrichment for H3K14ac in the mutant compared with WT (h). Values are means ± SD from three biological replicates. Each biological replicate corresponds to three technical replicates. Unpaired two-tailed Student’s t-test is employed to measure statistical significance between two samples (ns, no significance; **, *p* < 0.01; ***, *p* < 0.001).

### Elevated *AGO5* abundance in the nucellus disrupts the initiation of megagametogenesis

To test whether increased AGO5 abundance in the nucellus of pre-meiotic ovules might indeed causes defects in megagametogenesis, the *AGO5* coding sequence was cloned under the control of the MMC-specific *KNU* promoter or the broadly active nucellar *SWI3b* promoter and transformed into plants. Two of the three independent transgenic lines containing *pKNU:AGO5* exhibited fertilization rate comparable to those of the WT, whereas one line showed only a slight reduction. Consistent with these results, none of the three *pKNU:AGO5* lines displayed significant defects in female gametogenesis compared with the WT (Fig. 6a-d). In contrast, both independent *pSWI3b:AGO5* transgenic lines showed a marked reduction (∼50%) in fertilization rate and a corresponding increase (∼50%) in defects during the initiation of megagametogenesis. The affected ovules contained female gametophytes arrested at the one-nucleate stage (Fig. 6a-d). Collectively, these results indicate that SWI3b represses elevated *AGO5* abundance and probably other genes throughout the nucellus including the MMC. Lack of *AGO5* suppression disrupts the transition to megagametogenesis.

**Fig. 6.**
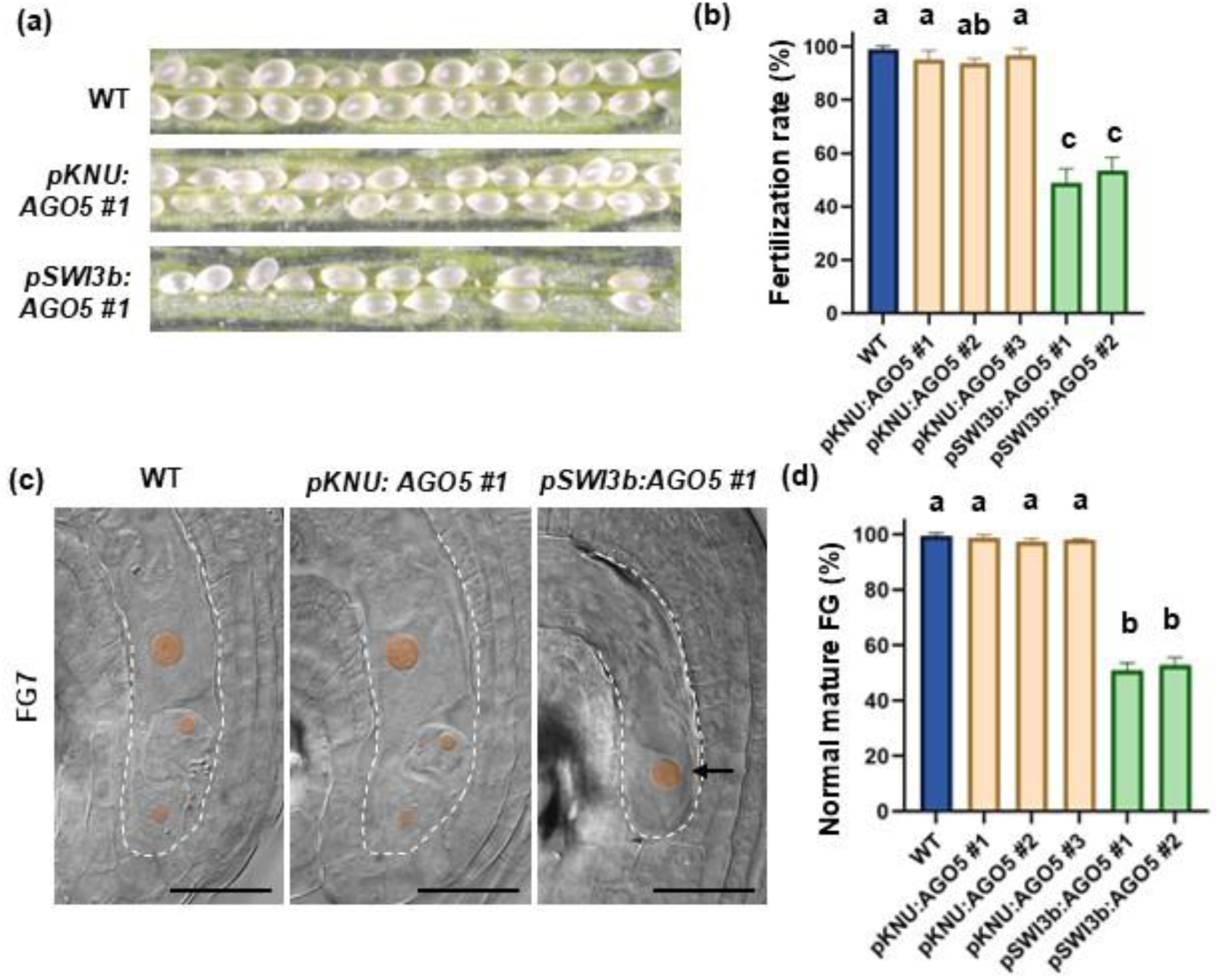
Elevated AGO5 abundance in the nucellus disrupts the initiation of megagametogenesis. (a) A representative dissected silique from WT, one transgenic line of *pKNU:AGO5* and one transgenic line of *pSWI3b:AGO5*. (b) Quantification of the fertilization rate from WT and indicated transgenic lines. Different letters above columns indicate significant differences (*P* < 0.05) as determined by one-way ANOVA. (c) DIC microscopy of ovules at FG7 in WT, one transgenic line of *pKNU:AGO5* and one transgenic line of *pSWI3b:AGO5*. Dashed line indicates the embryo sac. Nuclei in the female gametophyte are marked by red color. Bars, 20 μm. (d) Quantification of the percentage of WT-like female gametophyte in WT and indicated transgenic lines. Different letters above columns indicate significant differences (*P* < 0.05) as determined by one-way ANOVA.

## Discussion

Based on our findings, we propose that SWI3b functions as a chromatin-associated repressor that maintains *AGO5* transcription below a critical threshold in the nucellus before meiosis. We further suggest that SWI3b is recruited to the *AGO5* locus by one or more sequence-specific transcription factors, where it promotes histone deacetylation and limits H3K9 acetylation, plausibly by interacting with a histone deacetylase (HDAC). This repressive chromatin state prevents premature or ectopic AGO5 accumulation, thereby permitting the transition from meiosis to megagametogenesis. Loss of SWI3b disrupts this regulatory mechanism, leading to elevated H3K9ac of the *AGO5* locus, elevated *AGO5* expression in the nucellus, and failure to initiate the mitotic divisions of the functional megaspore. Together, our data reveals an epigenetic mechanism that coordinates chromatin remodeling with the developmental switch from meiosis to the initiation of megagametogenesis (Fig. 7).

**Fig. 7.**
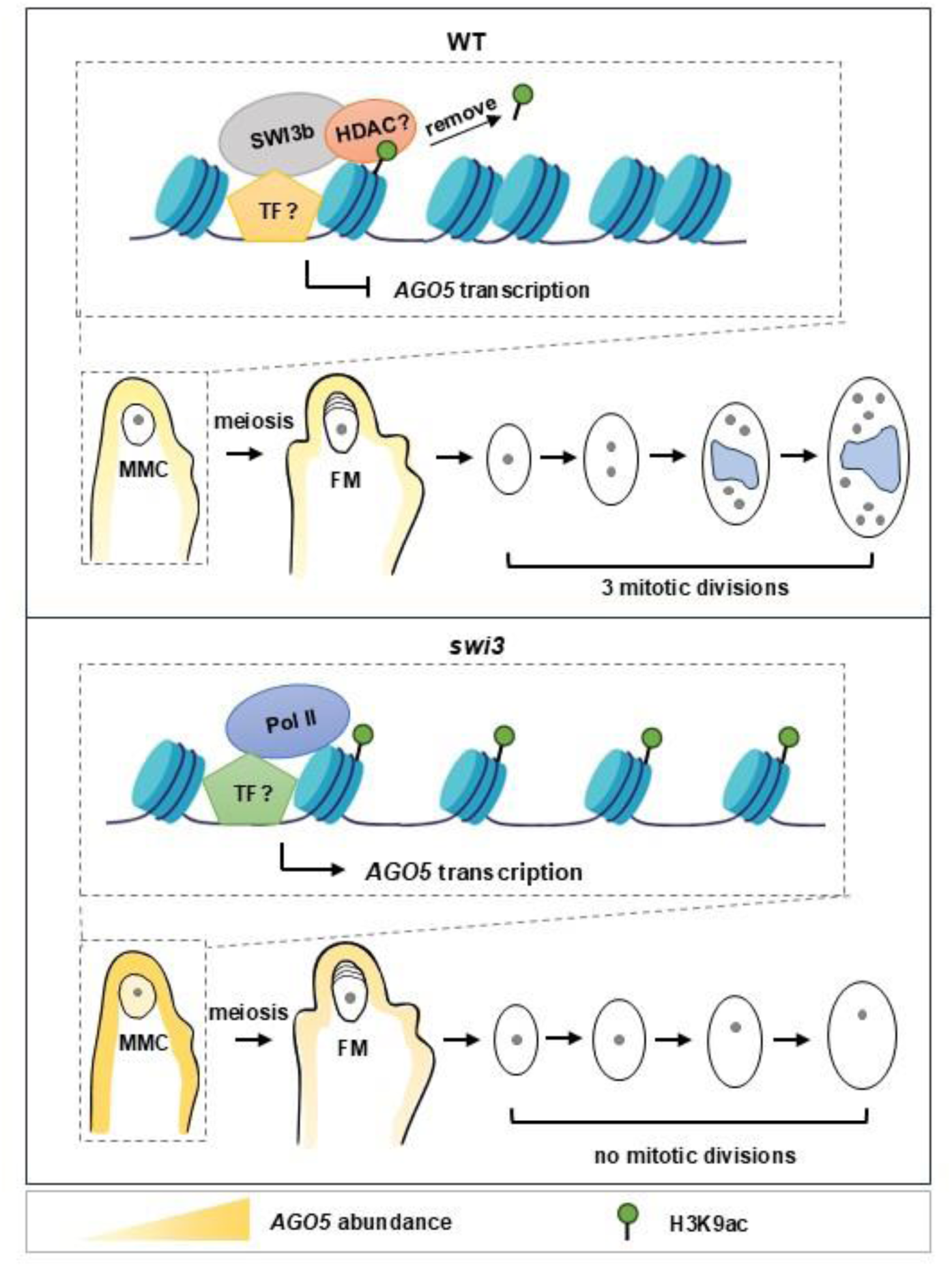
Model of SWI3b mediated *AGO5* expression to regulate the initiation of female megagametogenesis. In WT ovules, SWI3b is proposed to cooperate with an as-yet unidentified transcription factor (TF) to recruit histone deacetylase (HDAC) activity to the *AGO5* locus, thereby maintaining low H3K9 acetylation levels and repressing *AGO5* transcription before meiosis. Following meiosis, this repression is relieved, allowing the functional megaspore to initiate three rounds of mitotic divisions required for female gametophyte development. In the absence of SWI3b, increased H3K9 acetylation at the *AGO5* locus results in elevated *AGO5* expression in the nucellus. Our genetic and transgenic analyses indicate that ectopic AGO5 accumulation in nucellar cells, which interferes with the initiation of megagametogenesis, causing developmental arrest at the one-nucleate stage. The yellow to orange color gradient indicates the abundance of AGO5 protein. Green circles indicate H3K9ac marks. The as-yet unidentified factors are marked with question marks.

Many key developmental regulators of MMC formation and meiosis were identified. It is known that chromatin decondensation, small RNAs and epigenetic regulations contribute to MMC formation (Jiang & Zheng, 2021), but the factors controlling the transition to mitosis are not known. In *swi3* mutant ovules, the FM is formed but did not enter the first mitotic division. It fails most likely at cell cycle re-entry, or activation of S-phase genes or chromatin licensing of the mitotic program. SWI3b, as part of the MAS complex, may be required to switch from meiotic chromatin state to mitotic program. From analyzing the expression of cell cycle genes, we found that only *CDKA* is slightly downregulated in *ABI3 swi3*, but not *CDKB2;1*, *CDKB2;2*, *CYCB1;1*, *Histone 4* and *KNOLLE*. Since the defect occurs only in the functional megaspore, the effect of mis-regulation of gene expression might be diluted by qRT-PCR using the whole flower. The overall mitotic machinery may still operate relatively normal in most floral tissues, which is consistent with normal flower morphology and mature ovule morphology in *ABI3 swi3*. It is plausibly that *ABI3 swi3* affects *CDKA* expression in a small number of cells but does not strongly reduce the proportion of cells actively entering S phase, mitosis, or cytokinesis. CDKA is known to be required for mitotic divisions during male gametogenesis. In addition, CDKA and F-BOX-LIKE 17 (FBL17) control the first and second mitotic division cycles during megagametogenesis, which is consistent with SWI3b functioning in the same pathway (Dissmeyer *et al*., 2009; Zhao *et al*., 2012).

Previous studies implicated AGO5, a key effector of small RNA-mediated gene silencing in female reproductive development, is required for the initiation of megagametogenesis (Tucker *et al*., 2012). However, the mechanism underlying AGO5 regulation during this developmental transition has remained unclear. Our results demonstrate that SWI3b restricts *AGO5* expression in the nucellus by maintaining low H3K9ac at the *AGO5* locus. Notably, H3K9ac, but not H3K14ac, was increased in *ABI3 swi3-2*. This is consistent with previous findings that SWI3b and histone deacetylase 6 (HDA6) act together to reduce H3K9 acetylation, but not H3K14 or H3K27 acetylation, suggesting that the SWI3b–HDA6 complex exhibits specificity toward particular histone acetylation marks (Yang *et al*., 2020). SWI/SNF is a well-studied type of chromatin remodeling ATPase, which is mainly required for transcriptional activation. SWI3b belongs to the MINU-associated SWI/SNF complexes (Guo *et al*., 2022). It’s known in Arabidopsis, loss of function of MAS subunits (SWI3a, SWI3b, MINU1/2, SWP73, LFR, ARP7, PSA2) lead to lethality at reproductive stages, including female gemetogenesis (Kandasamy *et al*., 2005; Sarnowski *et al*., 2005; Sang *et al*., 2012; Sacharowski *et al*., 2015; Kong *et al*., 2020). It will now be interesting to examine whether other components from the MAS complex have similar functions in megagametogenesis like SWI3b.

Previous work demonstrated that complete loss of *AGO5* function has little or no effect on female gametophyte development, suggesting that the developmental defects associated with the semi-dominant *ago5-4* allele do not simply result from altered AGO5 activity, but rather from the acquisition of a novel function by the truncated AGO5 protein (Tucker *et al*., 2012). Tucker et al. proposed that AGO5-4 may sequester a subset of small RNAs or interfere with endogenous sRNA pathways, thereby disrupting signaling events required for the initiation of megagametogenesis. Our results identify an additional regulatory layer controlling AGO5 activity. We show that SWI3b restricts *AGO5* expression in the nucellus, likely through modulation of histone acetylation at the *AGO5* locus. These findings suggest that precise spatial restriction of AGO5 abundance, rather than simply its presence or absence, is essential for normal female gametophyte development. The requirement for tight regulation of *AGO5* expression is consistent with the broader importance of somatic small RNA pathways during ovule development. AGO9, for example, functions in nucellar companion cells to restrict the specification of reproductive cell fate through a non-cell-autonomous 24-nt siRNA pathway, illustrating that somatic tissues surrounding the female germline actively regulate reproductive development through epigenetic signaling (Olmedo-Monfil *et al*., 2010). Unlike AGO5, loss of AGO9 function only affects megasporogenesis. However, both proteins demonstrate that disruption of small RNA homeostasis in surrounding sporophytic tissues profoundly affects female gametophyte development. Our novel findings raise the possibility that SWI3b-dependent chromatin regulation maintains the appropriate balance of AGO-mediated small RNA activities in the nucellus during the transition from megasporogenesis to megagametogenesis. The detailed molecular nature of the AGO5-associated small RNA pathways in this process remains to be resolved in future studies. Because *ago5* null mutants exhibit no detectable defects in female gametophyte development, functional redundancy with other AGO proteins and their fine-tuned expression level is likely. Our data is compatible with this model and further suggest that chromatin-mediated repression of *AGO5* may prevent inappropriate activation or excessive accumulation of AGO5-associated silencing complexes within nucellar cells. In the *ago5-4* mutant allele, the truncated AGO5 protein may disrupt this pathway by aberrantly binding or sequestering specific classes of small RNAs, whereas in *swi3* mutants increased AGO5 abundance may similarly perturb the stoichiometry of AGO-containing silencing complexes. Thus, both, altered AGO5 activity and altered AGO5 dosage could interfere with the small RNA-mediated signaling required to initiate megagametogenesis.

Based on these observations, we propose a model in which SWI3b-dependent chromatin remodeling restricts *AGO5* expression in the nucellus by maintaining low H3K9 acetylation at the *AGO5* locus. This spatial repression ensures that AGO5 abundance remains below a threshold compatible with the proper transition to megagametogenesis. Loss of SWI3b function results in elevated *AGO5* expression, leading to disruption of small RNA-mediated regulatory pathways and inhibition of female gametophyte initiation. Future identification of the small RNAs associated with AGO5 and the factors responsible for recruiting SWI/SNF complexes to the AGO5 locus as well as further genomic loci under SWI3b control will be essential to further our understanding how chromatin regulation and small RNA pathways are integrated to control the initiation of female gametpgenesis in plants.

## Supplemental figures

**Fig. S1.**
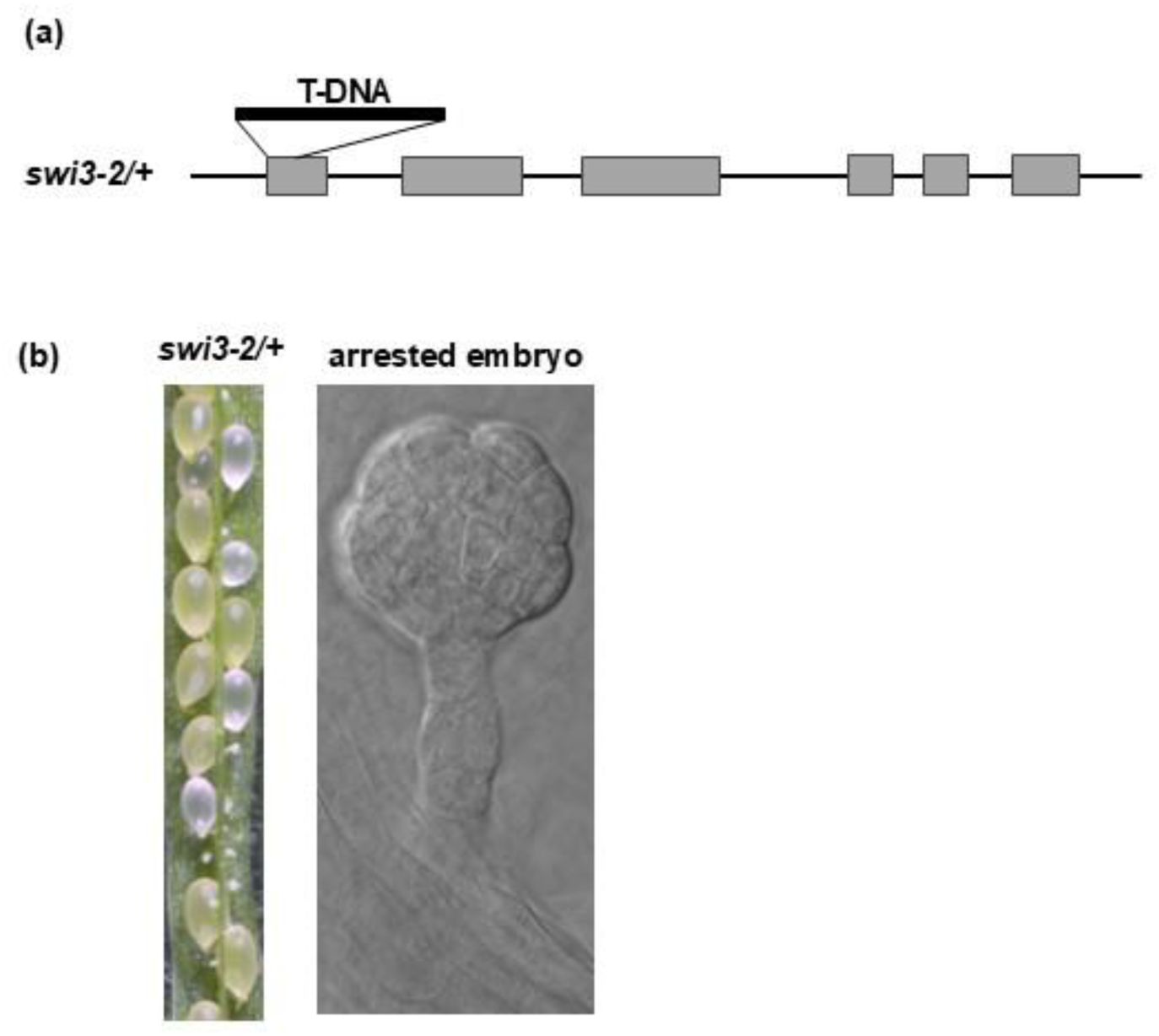
Embryo development of *swi3-2* was arrested at globular stage. (a) Schematic drawing showing *swi3b-2* (GABI_302G08) carrying a T-DNA in the first exon. (b) Dissected silique of *swi3-2/+* and DIC microscopy showing one arrested embryo selected from the whitish ovules.

**Fig. S2.**
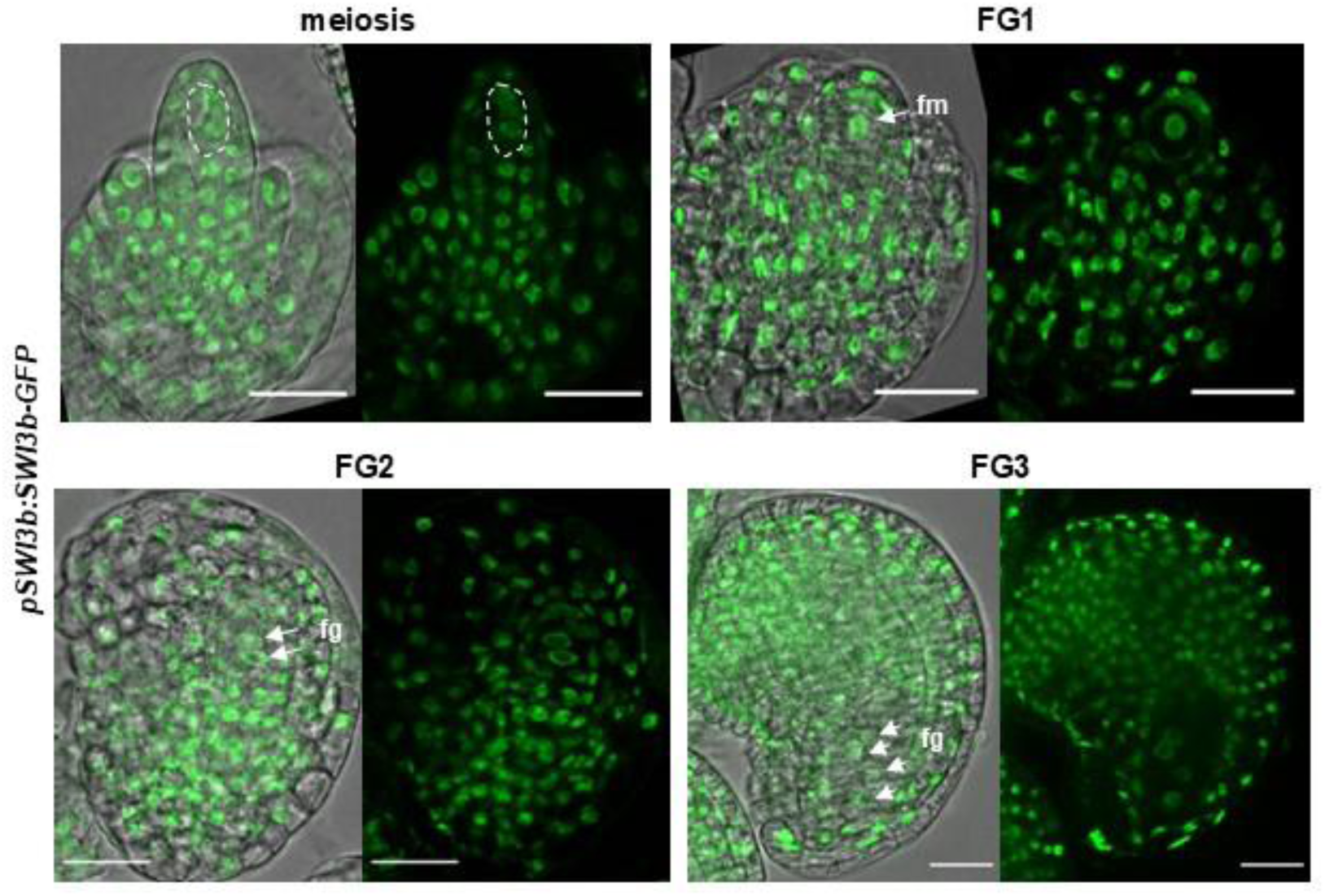
*SWI3b* is expressed throughout female gametogenesis. Expression pattern of *pSWI3b:SWI3b-GFP* in ovules undergoing meiosis (upper left), at FG1 (upper right), at FG2 (lower left) and at FG3 (lower right). Bars, 20 μm.

**Fig. S3.**
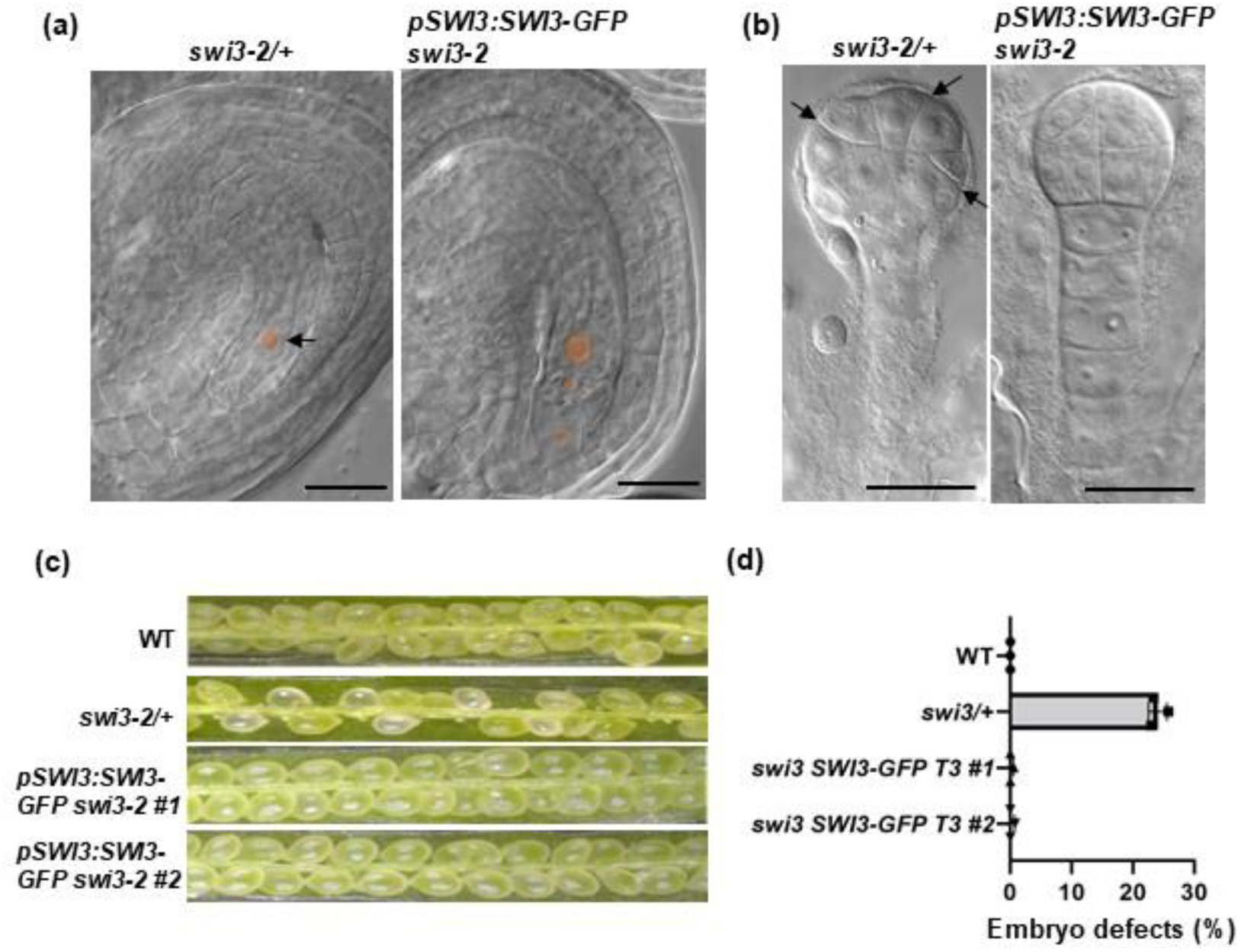
*pSWI3:SWI3-GFP* complemented the fertility defects of *swi3-2*. (a) DIC microscopy of ovules at FG7 in WT and one transgenic line of *pSWI3b:SWI3b-GFP* in a *swi3-2* homozygous background. Dashed line indicates the embryo sac. Nuclei in the female gametophyte are marked by red color. Bars, 20 μm. (b) DIC microscopy of 16-cell stage embryos in WT and *pSWI3b:SWI3b-GFP swi3-2*. Arrows point to abnormal cell division planes. Bars, 20 μm. (c) Dissected representative silique from WT, *swi3-2/+* and two independent transgenic lines of *pSWI3b:SWI3b-GFP* in *swi3-2*. (d) Quantification of the percentage of embryos arrested at the globular stage of the indicated genotypes.

**Fig. S4.**
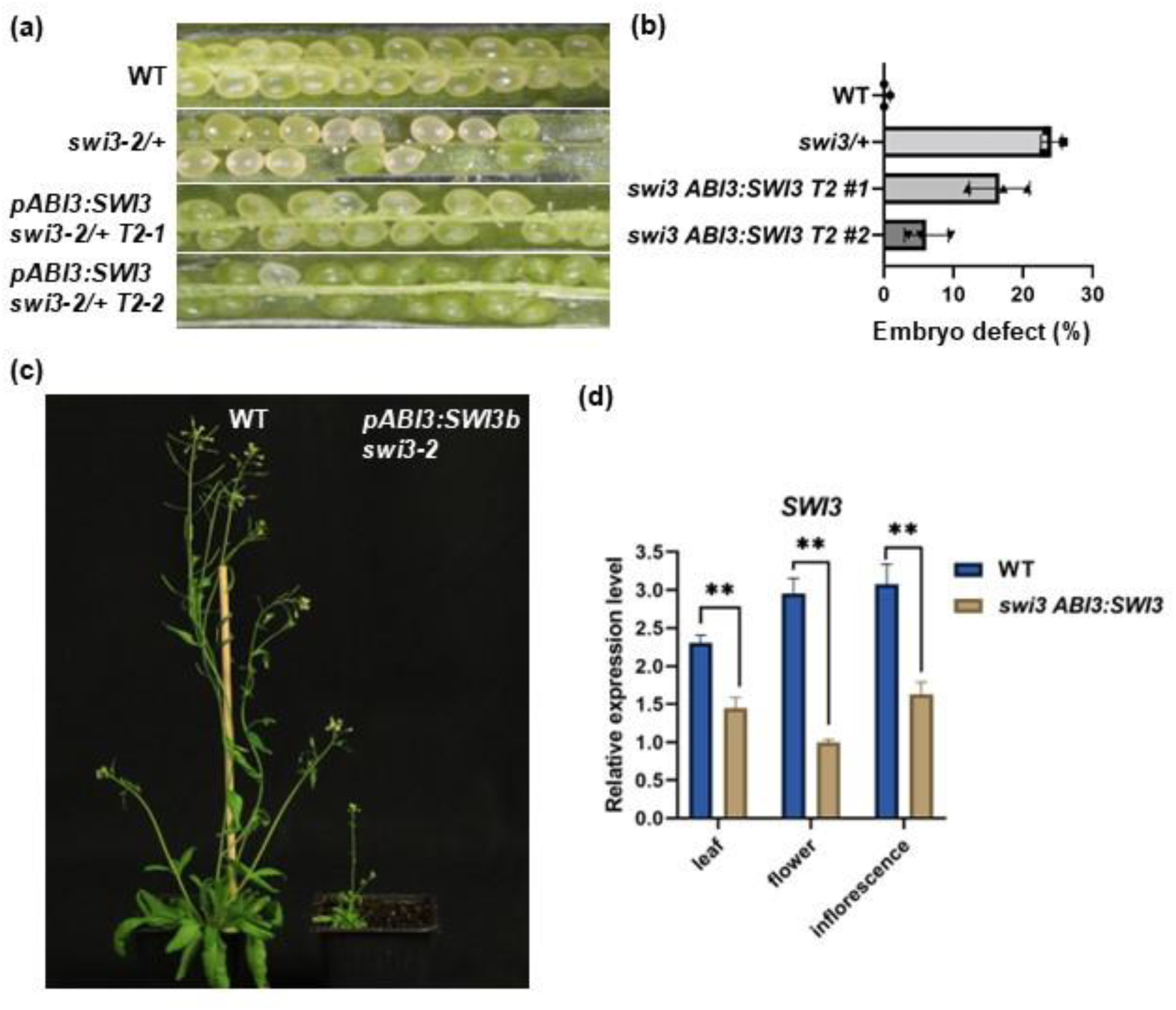
Embryo rescued *swi3-2* showed pleotropic developmental defects. (a) Dissected representative silique from WT, *swi3-2/+* and two independent transgenic lines of *pABI3:SWI3b-GFP* in *swi3-2/+*. (b) Quantification of the percentage of embryos arrested at the globular stage of indicated genotypes. (c) The flowering phenotype of a representative WT plant and one *pABI3:SWI3b swi3-2* homozygous plant. (d) Transcripts level of *SWI3b* in WT and *ABI3 swi3-2* leaf, flower and inflorescence, determined by quantitative RT-PCR. Values are means ± SD from three biological replicates. Each biological replicate corresponds to three technical replicates. Unpaired two-tailed Student’s t-test is employed to measure statistical significance between WT and the mutant (**, *p* < 0.01).

**Fig. S5.**
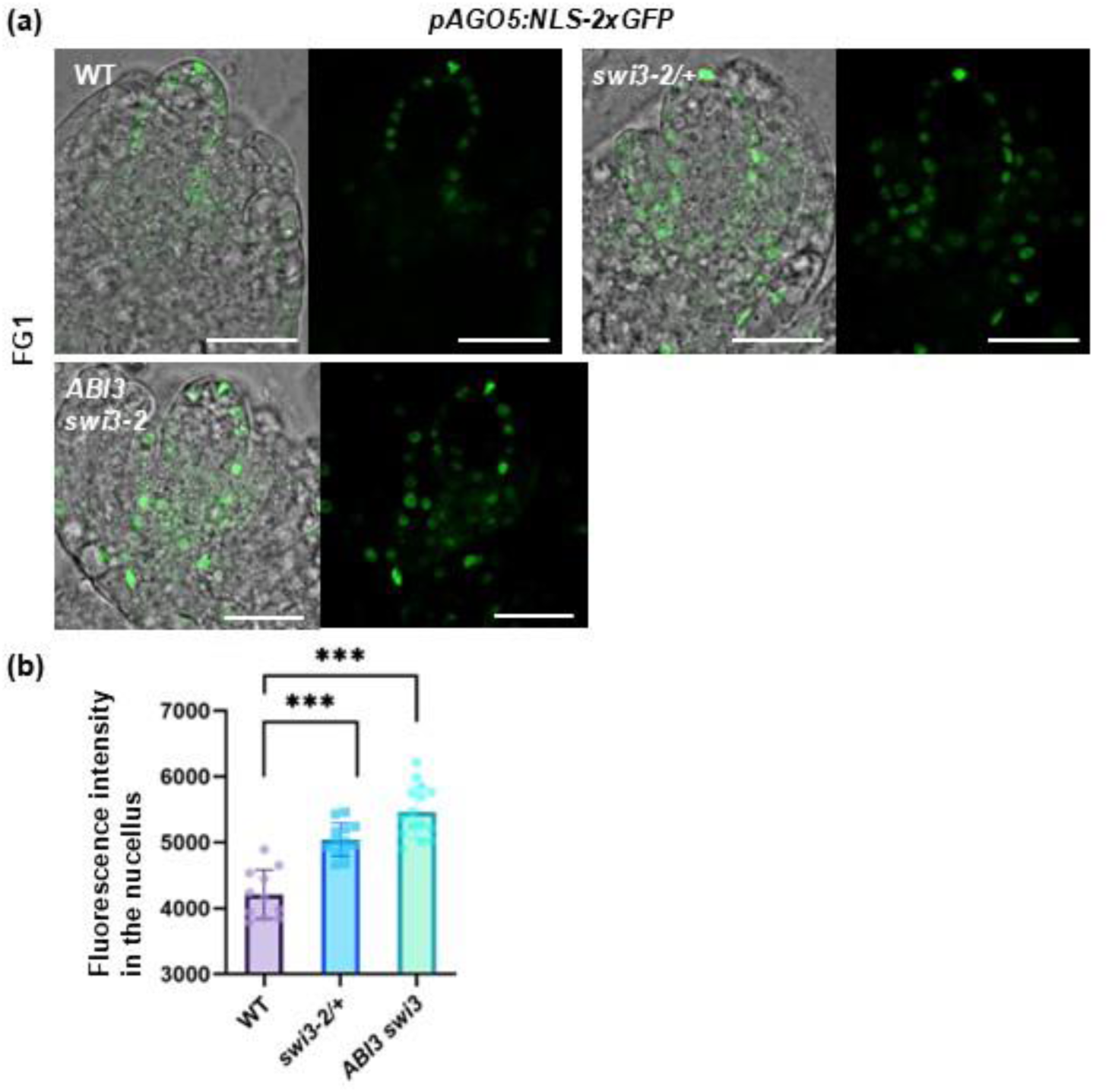
Expression pattern and strength of *AGO5* at FG1 was unchanged in *swi3*. (a) Expression pattern of *pAGO5:NLS-2xGFP* in WT, *swi3-2/+* and *ABI3 swi3-2* at FG1 stage. Bars, 20 μm. (b) Quantification of the average fluorescence intensity in the nucellus of indicated genotypes. One-way ANOVA followed by Dunnett’s multiple comparisons test, comparing each mutant line with the WT (ns, no significance).

**Fig. S6.**
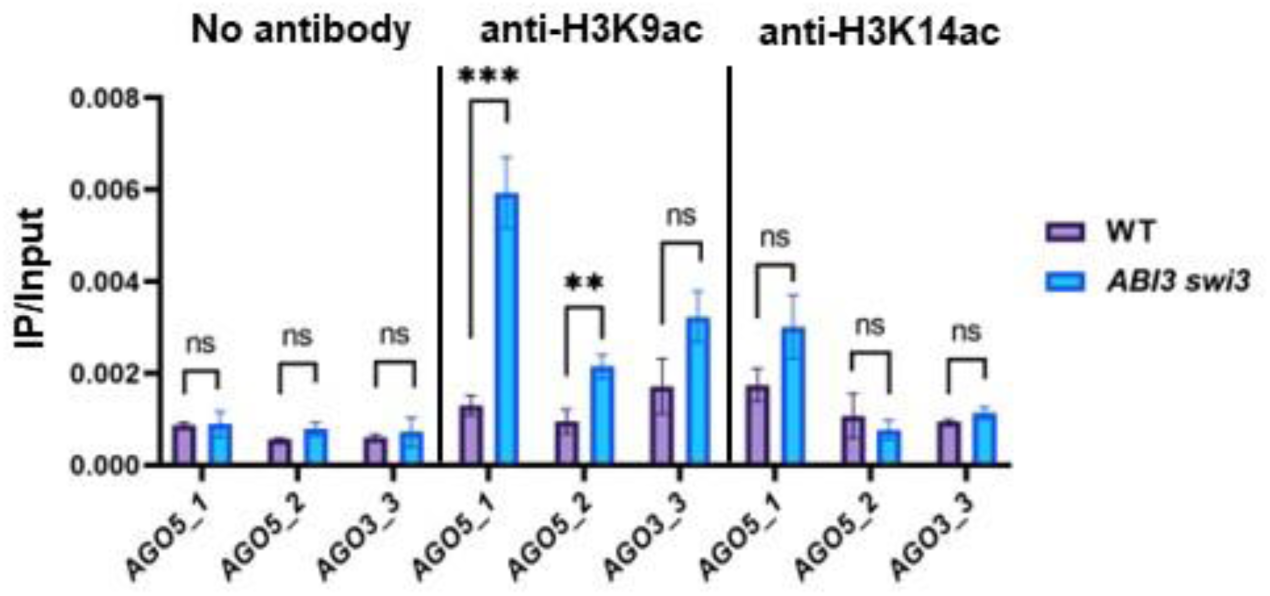
Enrichment of H3K9ac and H3K14ac in *AGO5* genomic regions. ChIP-qPCR analysis showing lack of enrichment for no antibody, enrichment for H3K9ac in region 1 and 2 of *AGO5* in *ABI3 swi3-2* inflorescence compared with WT, and lack of enrichment for H3K14ac in the mutant compared with WT. Values present the ratio of the relative quantity in the ChIP sample to that in the input sample. Values are means ± SD from three biological replicates. Each biological replicate corresponds to three technical replicates. Unpaired two-tailed Student’s t-test is employed to measure statistical significance between two samples (ns, no significance; **, *p* < 0.01; ***, *p* < 0.001).

## Acknowledgements

We thank Armin Hildebrand for plant care, Maria Hutterer for the lab work, and Tomasz Sarnowski, who kindly provided us the mutant seeds of *swi3-2/+*. The German Research Foundation (DFG) is acknowledged for financial support via SFB960 (to TD).

## Competing interests

None declared.

## Author contributions

WG initiated and designed the research. WG performed experiments. LL and HC provided transgenic reporter lines. TD provided research funding and input. WG and TD wrote the manuscript with input from all authors.

## Data availability

All data are available in the manuscript or the Supporting Information.

## Notes

### Competing Interest Statement

The authors have declared no competing interest.

